# A scalable random walk with restart on heterogeneous networks with Apache Spark for ranking disease-causing genes using type-2 fuzzy data fusion

**DOI:** 10.1101/844159

**Authors:** Mehdi Joodaki, Nasser Ghadiri, Zeinab Maleki, Maryam Lotfi Shahreza

**Affiliations:** Department of Electrical and Computer Engineering, Isfahan University of Technology, Isfahan 84156-83111, Iran

**Keywords:** Prioritization, Type-II Fuzzy Voter, RWRHN, Genes associated with the disease, Gene-Gene network, Heterogeneous networks

## Abstract

Prediction and discovery of disease-causing genes are among the main missions of biology and medicine. In recent years, researchers have developed several methods based on gene/protein networks for the detection of causative genes. However, because of the presence of false positives in these networks, the results of these methods often lack accuracy and reliability. This problem can be solved by using multiple genomic sources to reduce noise in data. However, network integration can also affect the quality of the integrated network. In this paper, we present a method named RWRHN (random walk with restart on a heterogeneous network) with fuzzy fusion or RWRHN-FF. In this method, first, four gene-gene similarity networks are constructed based on different genomic sources and then integrated using the type-II fuzzy voter scheme. The resulting gene-gene network is then linked to a disease-disease similarity network, which itself is constructed by the integration of four sources, through a two-part disease-gene network. The product of this process is a reliable heterogeneous network, which is analyzed by the RWRHN algorithm. The results of the analysis with the leave-one-out cross-validation method show that RWRHN-FF outperforms both RWRHN and RWRH. The proposed method is used to predict new genes for prostate, breast, gastric and colon cancers. To reduce the algorithm run time, Apache Spark is used as a platform for parallel execution of the RWRHN algorithm on heterogeneous networks. In the test conducted on heterogeneous networks of different sizes, this solution results in faster convergence than other non-distributed modes of implementations.

## 1. Introduction

Empirical identification and ranking of disease-causing genes are both time-consuming and expensive. However, finding these causative genes can have a high impact on disease prevention, treatment, and drug development and offer insights into gene functions and pathways. Therefore, understanding the relationship between diseases and genes is a crucial role expected from bioinformatics.

A common way of ranking disease-causing genes is the network-based approach (X. Wang, Gulbahce, & Yu, 2011). It aims at comparing candidate genes with known disease-related genes and rate them accordingly. A common method that is used in the current literature in the network-based approaches is based on considering local measurement. A widely used local measurement method is the one proposed by Dezso et al. (Dezső et al., 2009), which involves using a local measure called the shortest path betweenness to find the causative genes. In (Zeng, Zhang, & Zou, 2015), researchers examined the miRNA network to identify miRNAs that contribute to diseases through dysfunction or dysregulation. Xue et al. (X. Jiang, Zhang, Quan, Liu, & Yin, 2017) labeled the gene expression network with a propagation algorithm to discover the clusters associated with disease-causing genes. Sipko et al. (van Dam, Vosa, van der Graaf, Franke, & de Magalhaes, 2017) investigated various tools and methods of analysis of the co-expression network with RNA sequencing for the discovery of disease-causing genes. In [1], the author proposed to consider hubs in PPI or gene-gene networks, based on the fact that causative genes of some diseases form a hub. However, this method only works for some of the diseases. Indeed, disease-causing genes are not always directly linked together. DIGNiFI (Liu, Yang, Lin, Simmons, & Lu, 2017) uses a combination of global and local measures on the PPI network to find disease-causing genes, we will shortly explain more global methods.

The network-based approaches are highly dependent on the quality and accuracy of biometric networks. In other words, the fewer the false-positives (Montañez & Cho, 2013) in the interactions networks, the more reliable will be the results. The network-based approaches typically combine different genomic networks such as Gene Ontology (GO) (Schlicker, Lengauer, & Albrecht, 2010), gene expression (van Dam et al., 2017), protein sequences (Adie, Adams, Evans, Porteous, & Pickard, 2005), and Protein-Protein Interaction (PPI) networks from Human Protein Reference Database (HPRD) (Peri et al., 2003) for ranking disease-causing genes.

Moreover, limiting the ranking to just one type of network, such as genomic networks, may not always produce a reasonable estimate of causative genes genomic networks. It is known that biologically similar genes are more likely to influence a particular disease or a group of similar diseases (Ideker & Sharan, 2008; Oti & Brunner, 2007). Therefore, it is common to use a combination of genomic networks and disease similarity networks to increase the accuracy of detecting disease-causing genes. HeteSim (Zeng, Liao, Liu, & Zou, 2017), combines the disease similarity networks derived from MimMiner (Van Driel, Bruggeman, Vriend, Brunner, & Leunissen, 2006) and the PPI network from HPRD (Peri et al., 2003) and the Human Net to construct a heterogeneous network. Then it scores the gene-disease paths accordingly. In the method called GLADIATOR (Silberberg, Kupiec, & Sharan, 2017), disease-causing protein modules are predicted by combining the knowledge derived from the disease similarity network and disease-related proteins.

Since not all disease-causing genes are directly connected, recent studies have tried to use algorithms that would cover the entire network (Y. Li & Patra, 2010; Liu et al., 2017). Any network resulting from the integration of disease and gene networks would consist of nodes of different types. Considering this fact, Liu et al. (Y. Li & Patra, 2010) have modified the Random Walk with Restart (RWR) algorithm into a new algorithm called Random Walk with Restart on Heterogeneous Network (RWRHN). The RWRHN algorithm conducts a global search on the entire network, which produces better results than local methods. In several recent studies such as (Y. Li & Patra, 2010; Luo & Liang, 2015; Stenson et al., 2009; Tian et al., 2017), candidate genes have been ranked by creating a different heterogeneous network and running RWRHN on the entire network.

Jiawei et al. (Luo & Liang, 2015) analyzed the correlations between protein pairs and reconstructed the PPI network accordingly to remove false interactions from this network. They then used the RWR algorithm to rank candidate genes on the network. In a study by Le et al. (Le & Dang, 2016), data from different sources were integrated to construct a more reliable heterogeneous network. Specifically, Human Phenotype Ontology (HPO) (Köhler et al., 2013) and Online Mendelian Inheritance in Man (OMIM) (OMOM, 2014) database (disease similarity network based on text mining) were used to improve the disease interaction network. Also, the gene expression data, GO, and PPI was used to improve the accuracy of the gene-gene interaction network.

A serious weakness of existing network-based methods is their sensitivity to the noise. Since the presence of noise in the biological networks is unavoidable, the problem of noise and false interactions is best to be handled by integrating the networks derived from different bio-data sources (Lee, Blom, Wang, Shim, & Marcotte, 2011; Y. Li & Patra, 2010; Luo & Liang, 2015; Mehranfar, Ghadiri, Kouhsar, & Golshani, 2017; Tian et al., 2017; Zeng et al., 2017). However, a challenging task in integrating different data sources, is the method of this integration that highly affects the quality of the results (Mehranfar et al., 2017).

The main contributions of the present study are (1) constructing a more enriched network, (2) proposing a more accurate integration method, and (3) offering a highly efficient parallel RWRHN algorithm for the proposed method based on Spark. We constructed the gene-gene and disease-disease interaction networks from different sources and then integrated these networks through a two-part gene-disease network to ultimately construct a reliable heterogeneous network.

To build a reliable gene-gene network, sources were combined using the type-II fuzzy voter scheme (Karnik & Mendel, 2011) introduced by Mehranfar et al. (Mehranfar et al., 2017). To discover protein complexes, Mehranfar et al. first used interval type-II fuzzy voter scheme to reduce noise in the integration of GO and gene expression data networks. Then they analyzed the resulting network in the search for protein complexes.

In this study, a fuzzy method was used in place of classical integration methods to reduce the rate of false positives in different sources. A significant advantage of this fuzzy approach over conventional methods is that the weights are given to each interaction in the network rather than the entire source. Considering the large size of the resulting heterogeneous network and the inability of RWRHN to finish a global search over the whole network in a reasonable time, the Apache Spark platform was used for parallel execution of this algorithm.

In this paper, we will present the developed method, called RWRHN with Fuzzy Fusion (RWRHN-FF), for building a reliable heterogeneous network by integrating multiple networks derived from different sources with the help of Type-II fuzzy voter scheme. In Section 5.2, the predicted genes of RWRHN-FF method on the heterogeneous networks constructed for prostate, breast, gastric, and colon cancers will be presented as evidence of performance. The comparisons between the results of RWRHN-FF, RWRH (Y. Li & Patra, 2010), and RWRHN (Luo & Liang, 2015) in terms of AUC and Precision will show that RWRHN-FF outperforms both of its predecessors. It will also be shown that the gene-gene network constructed with the type-II fuzzy voter scheme provides better estimates than the one constructed by the classical (averaging) method.

## 2. Methods

The paper will also explain how the Apache Spark platform (*Graphx: A resilient distributed graph system on spark*, spark) is used for parallel execution of the algorithm to achieve shorter run times.

### 2.1 Datasets

This section provides a brief explanation about the sources and methods used for constructing individual networks that were ultimately compiled into the heterogeneous network (all steps shown in Fig.1.).

**Fig. 1.**
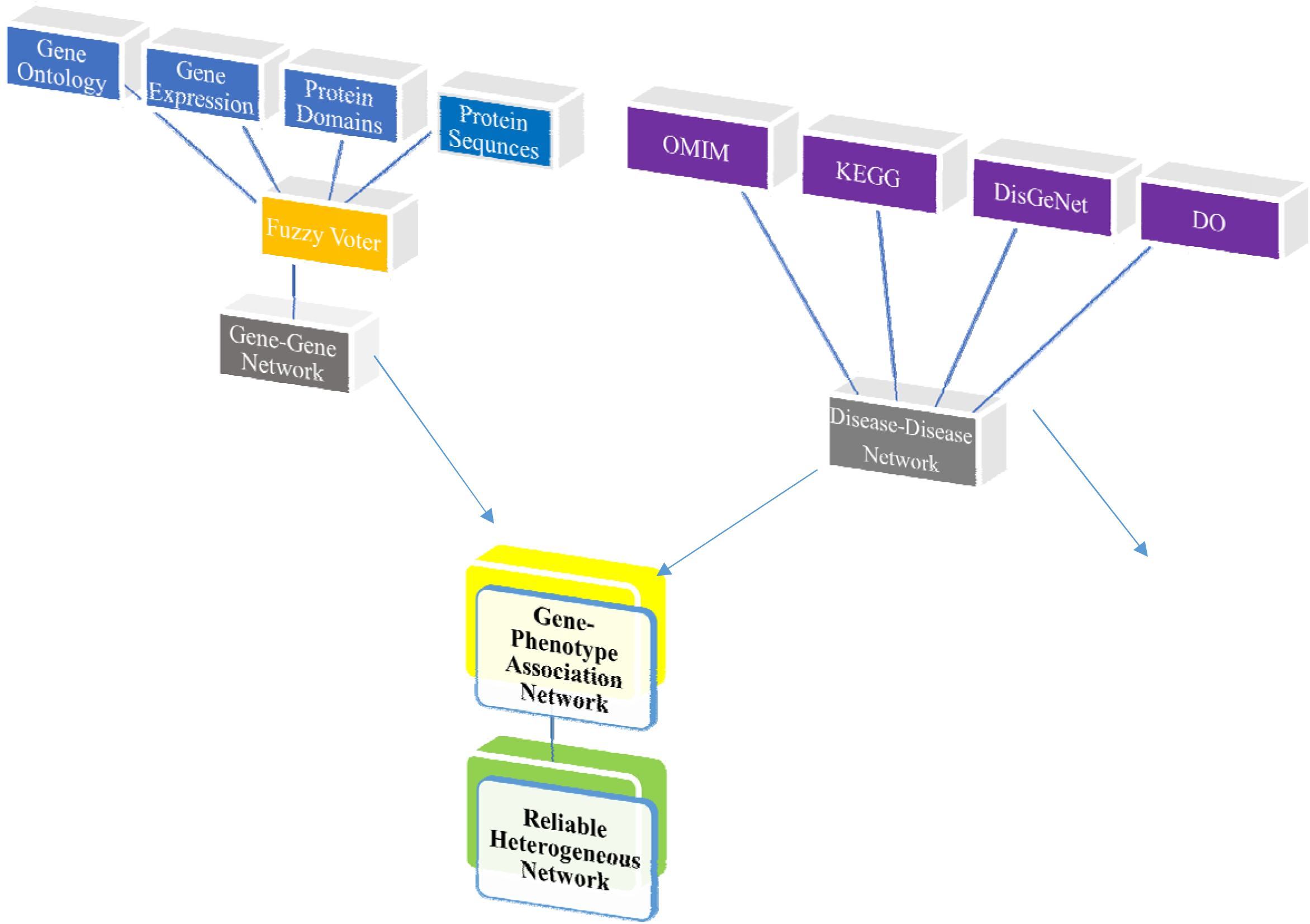
The construction process of reliable gene-gene network.

#### 2.1.1 Gene-gene interaction network

this network was constructed based on the data of four sources. After building a gene-gene sub-network for each source, the four sub-networks were integrated using the type-II fuzzy voter scheme.

The first gene-gene sub-network was constructed based on Gene Ontology Annotations (Schlicker et al., 2010), namely cellular component (CC), molecular function (MF) and biological process (BP) using the method of Wang (J. Li et al., 2011; J. Z. Wang, Du, Payattakool, Yu, & Chen, 2007) for 20,083 genes. After analyzing the descriptions of each gene and its related products using CC, MF and BP ontologies, the similarities of gene pairs were collated into a matrix.

The second gene-gene similarity sub-network was constructed based on the protein sequences in the Uniprot (Ashkenazy, Erez, Martz, Pupko, & Ben-Tal, 2010) database using the Basic Local Alignment Search Tool (BLAST) (Altschul et al., 1990). For this sub-network, the similarity of 20,412 proteins was calculated by comparing the strands of protein sequences with a threshold value of 10^−6^. Then the BIT SCORE of similarity between proteins was normalized by the MIN-MAX method.

The third gene-gene sub-network was constructed based on COXPRESSdb (Obayashi & Kinoshita, 2010) database using the mutual rank measure. Since a gene may not behave the same in different tissues and at different times, it is necessary to analyze COXPRESSdb with the mutual rank to measure the behavioral similarity of genes in various tissues.

The fourth sub-network was constructed based on the similarity of protein domains for 18,000 proteins in the Pfam (Bateman et al., 2011) database (within its protein sequence, each protein has a set of sub-sequence that determine the function and family of that protein). For this sub-network, the similarity between each pair of proteins was computed based on the families to which they belong using the Jaccard’s method (Jaccard, 1908).

#### 2.1.2 Phenotype similarity network

another essential network for the discovery of disease-causing genes is the disease similarity network. Like the gene-gene similarity network, which can be made from different sources, the disease similarity network can be constructed based on various criteria such as shared disease gene, shared pathways, and shared disease ontology.

Following the approach of the Heter-LP algorithm for drug repositioning (Shahreza, Ghadiri, Mousavi, Varshosaz, & Green, 2017), we used four sources to develop the disease similarity network with better weighting of disease-disease interactions: 3,321 cases of similarity in terms of causative genes according to DisGeNet (Stenson et al., 2009); 4,798 cases of phenotypic similarity according to OMIM (OMOM, 2014) database, 9,679 cases of phenotypic similarity according to DOSE (Osborne et al., 2009) database, and 1,366 cases of similarity in terms of biological pathways and causative genes according to KEGG (Bock & Goode, 2002) database. After measuring the similarity between each pair of diseases in each source, a similarity matrix was constructed for each source separately and the four obtained matrices were integrated to construct the disease-disease similarity network.

#### 2.1.3 Gene-phenotype association network

To create a heterogeneous network, it was necessary to develop a two-part gene-disease network. In other words, each of the sources (OMIM, KEGG, DO and DisGeNet) contains data about diseases and about disease-causing genes, which had to be integrated to form a two-part gene-disease network. Ultimately, this network was used to link the gene-gene network to the disease-disease network and construct a reliable heterogeneous network.

### 2.2 The hybrid method for reducing the impact of false interactions on the gene-gene network

Since each source assigns a certain weight to the interaction between every two genes, the interaction weights considered in each source can influence the fusion of sources and the final network. In this study, instead of giving an overall weight to each source, we used the type-II fuzzy voter scheme to give a weight to each interaction. There are several ways to integrate biological data. While simple methods such as averaging are easy and straightforward, the methods based on fuzzy logic are more suitable for this purpose. A key feature of fuzzy logic is that it allows us to give each source a degree of accuracy. With this feature, it is possible to lower the impact of interactions that have been generated incorrectly as a result of experimental (Mehranfar et al., 2017). Further, giving an overall weight to a source can never be an effective approach to address all its shortcomings. The general structure of the fuzzy inference system is illustrated in Fig**. 2**. As shown in this figure, the fuzzy knowledge base of the system is constructed by a set of if-then rules. The fuzzifier component uses the membership function in the fuzzy knowledge base to convert numeric inputs into linguistic (fuzzy) variables. The inference engine uses the if-then rules to convert fuzzy inputs into fuzzy outputs. And finally, the defuzzifier component uses the membership functions to convert the output of the fuzzy inference engine to numerical outputs.

**Fig. 2.**
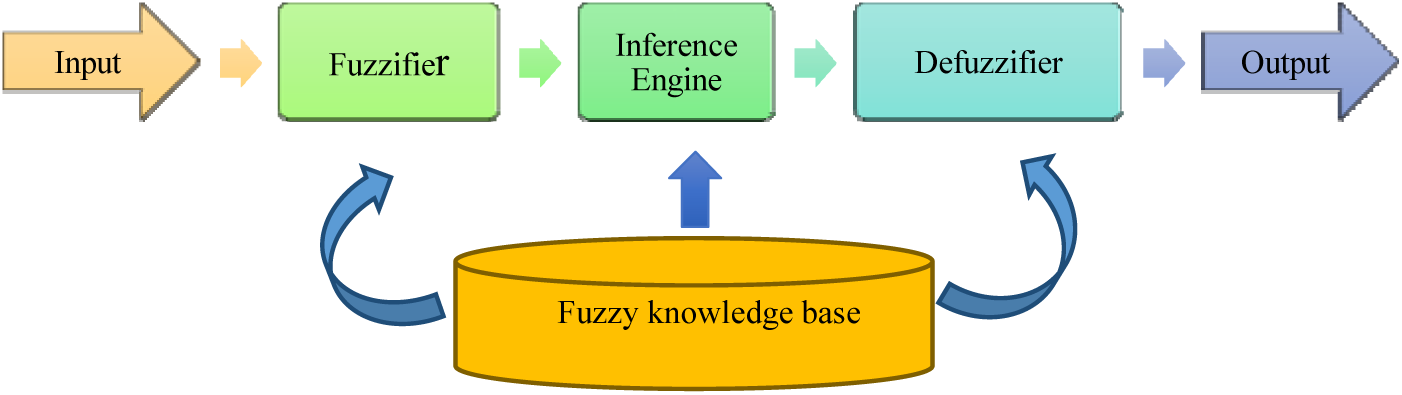
The architecture of the fuzzy inference.

#### 2.2.1 Type II fuzzy system

In type-I fuzzy systems, the membership functions are determined by an expert. However, this opinion is subject to change, and also different experts may have different views about the shape of the membership function. In other words, it is possible to encounter uncertainty or a lack of consensus regarding the boundaries of the membership function. Type-I fuzzy logic cannot model these uncertainties. But, type-II fuzzy logic can be used to resolve this problem (Linda & Manic, 2011). In this study, Gaussian membership functions were considered to be in the form of intervals. By addressing the said uncertainty in type-I fuzzy logic, type-II fuzzy logic can help us achieve a more flexible model of the studied phenomenon. In many ways, type-II fuzzy logic can be seen as an extended version of type-I. Whereas in type-I fuzzy logic, the membership degree is just a number, in type-II, this degree itself constitutes a fuzzy system. For problems where there are some uncertainties in membership functions, it is better to use type-II fuzzy logic instead of type-I (for more detailed information about the structure and function of the Type-II fuzzy voter scheme, see (Linda & Manic, 2011)).

Here, we use the definitions given in (Mehranfar et al., 2017) for fuzzy rules and membership functions. As shown in Fig.5, the four inputs *x*_*1*_, *x*_*2*_, *x*_*3*_, and *x*_*4*_ are assumed to represent the weights of the interaction between genes *g’* and *g”* in sources *S*_*1*_, *S*_*2*_, *S*_*3*_, and *S*_*4*_, respectively. To compute a weight for the *g’*-*g”* interaction in *S*_*1*_ with the type-II fuzzy voter scheme, the first step is to calculate the difference between the weight given to this interaction in *S*_*1*_ and that in other sources (*d*_*ij*_*=|x*_*i*_-*x*_*j*_*|*). Then, the membership functions named Small, Medium, and Large (Fig.3) are used to fuzzify these differences (convert the numerical inputs into fuzzy sets). Depending on the defined range of membership functions, an input may become a member of one or multiple functions. The next step is to use the fuzzy inference rules and the membership functions (Vlow, Low, Medium, High, and Vhigh shown in Fig.4) to map the fuzzy inputs to the fuzzy outputs, or in other words, compute the Agreeability of the source *S*_*1*_ with other sources. For example, in Eq.1, if *d*_*12*_ and *d*_*13*_ are members of the Small function and *d*_*14*_ is a member of the Medium function, then the output will be Vhigh. Some of the fuzzy rules used in this study are listed in Table.1. Next, the type-II fuzzy set needs to be reduced to type-I. In the present study, this conversion is performed using the method of Karnik and Mendel (Karnik & Mendel, 1998). The final stage is defuzzification or computing the final weight for the *g’*-*g”* interaction in *S*_*1*_.

**Table.1.**
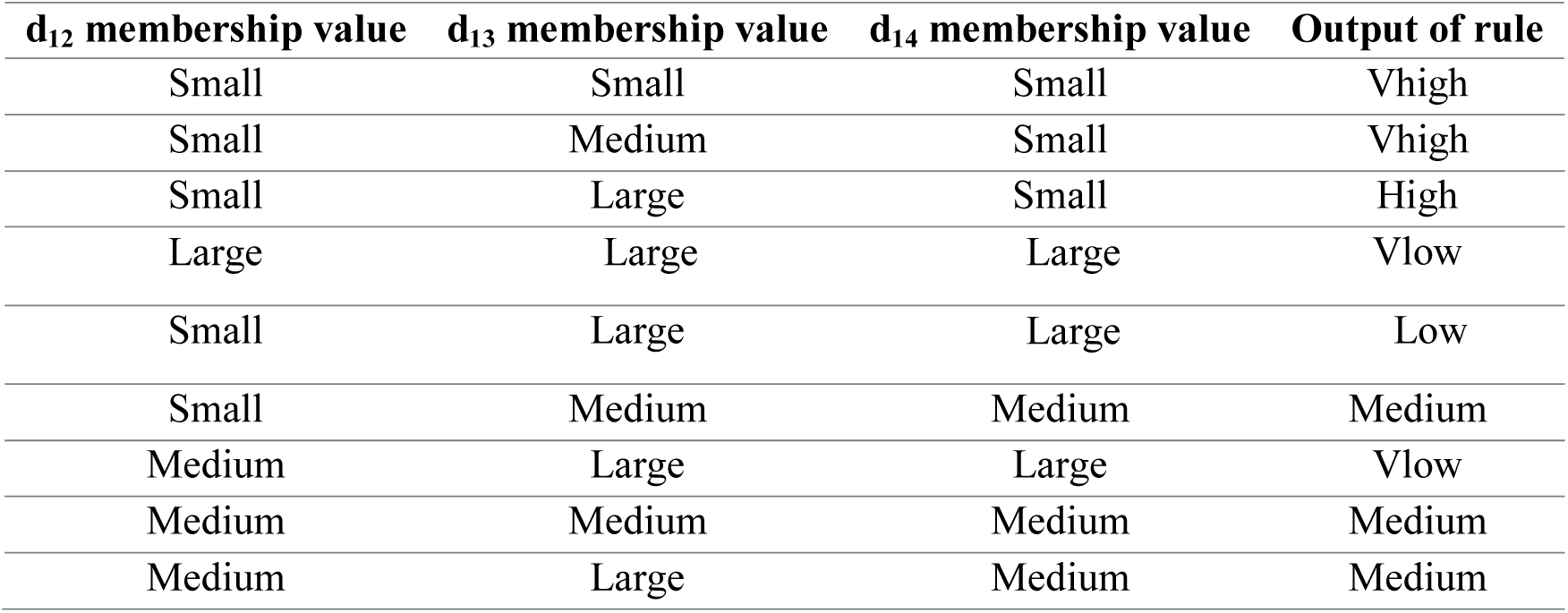
Some of linguistic rules.

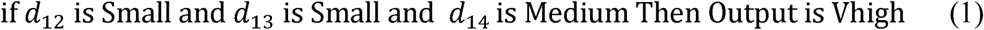

**Fig.3.**
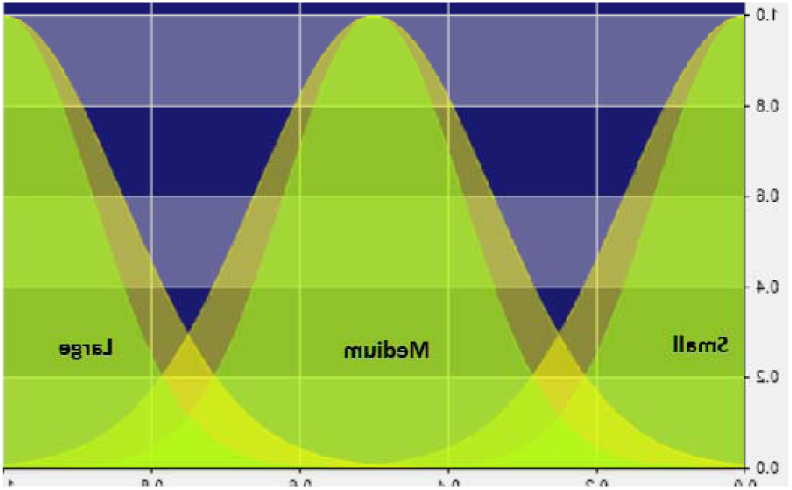
Membership function for fuzzification.

**Fig.4.**
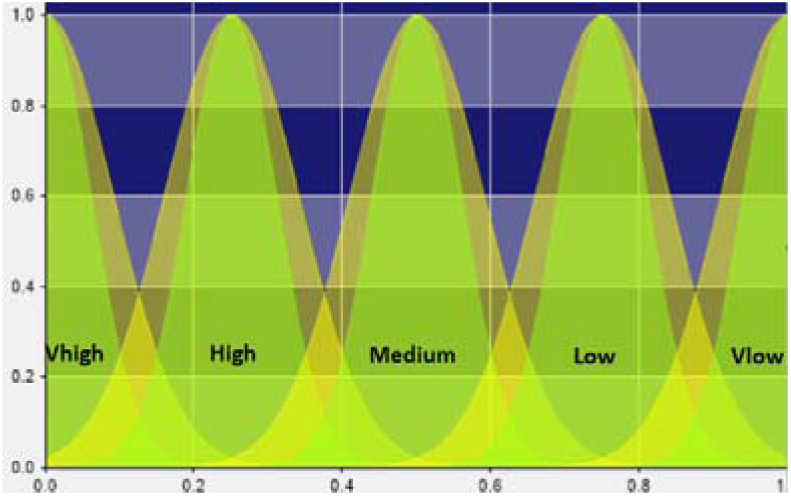
Membership function for determing fuzzy agreeability.

**Fig.5.**
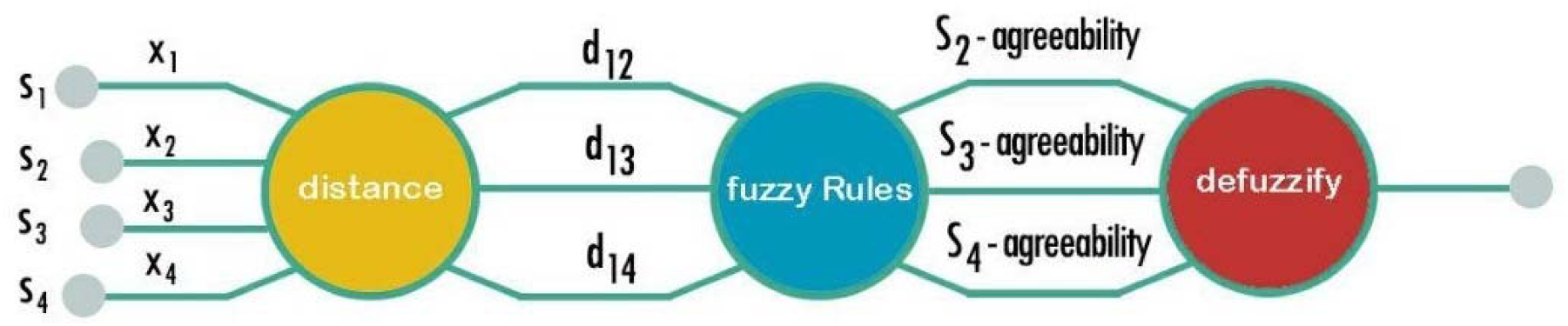
The architecture of the fuzzy voter.

After computing the Centroid *C*_*j*_ of the outputs of the sets *B*_*j*_, the weight source of 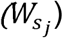 for *S*_*1*_ 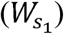 is obtained from Eq.2. In this equation, *C*_*j*_ is the Centroid of the fuzzy sets *B*_*j*_ derived from the rules, 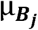 is the membership value of the inputs of the fuzzy sets *B*_*j*_, and *M* is the number of fuzzy sets.

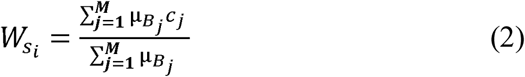

After computing the weight of the *g’*-*g”* interaction in each source as described above, the final weight of this interaction must be obtained using Eq.3. In this equation, 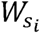 is the weight given by the fuzzy voter to the source *i* and *x*_*i*_ is the initial weight of the interaction between the two genes in the source *i*.

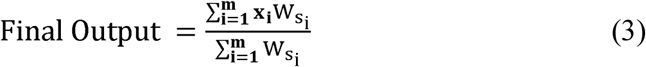

## 3. Ranking of candidate genes

RWRH (Y. Li & Patra, 2010) algorithm runs a global search on the networks, which provides a better ranking of the nodes and therefore a better prediction of disease-causing genes than local methods. This algorithm starts with a set of seed nodes and searches the entire network for similar nodes until convergence. The mathematical formulation of the RWRHN algorithm is provided by Eq.4:

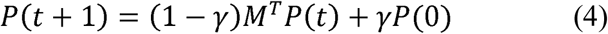

where, *P(t+1)* denotes the rank of the network nodes in step *t+1*, and parameter *γ* is a value between 0 and 1 which determines the probability of the jumping to seed nodes in the vector *P(0)*. Before executing the algorithm, we construct the matrix *M*, which consists of disease-disease, gene-gene, disease-gene, and gene-disease networks:

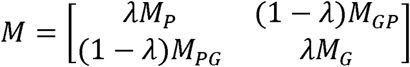

In the above matrix, *M*_*G*_, *M*_*P*_, and *M*_*GP*_ denote the gene-gene network, the disease-disease network, and the two-part disease-gene network, respectively. The parameter *λ* is the probability of jumping from the gene-gene network to the disease-disease network and vice versa. The vector *P(0)* is as follows:

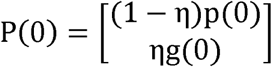

In the above vector, *P(0)* and *g(0)* are the initial probability of seed nodes and *η* denotes the importance of disease and gene seed nodes. After several iterations, the algorithm converge. The probability of *P(∞)* will be:

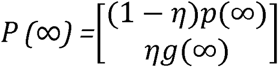

### 3.1 Parameter setting for RWRHN

According to the above description of RWRH, to discover disease-causing genes with this algorithm, we specify three parameters *γ, λ*, and *η*. After reviewing the previous works in this area such as (Le & Dang, 2016; Y. Li & Patra, 2010; Luo & Liang, 2015; Osborne et al., 2009; Tian et al., 2017), it was found that the best values for *γ, λ*, and *η* would be 0.7, 0.8, and 0.5, respectively.

Using more bio-data in the construction of gene-gene and disease-disease networks to reduce source noise inevitably increases the size of the final heterogeneous network. Given the global nature of RWRHN (that searches the entire network), it takes a long time for this algorithm to converge on such heterogeneous networks. In other words, using more sources will dramatically increase the number of network nodes and their interactions, which will impose a massive computation load on RWRHN at every iteration.

### 3.2 Execution of RWRHN on Apache Spark platform

Apache Spark uses the concept of RDD (*Graphx: A resilient distributed graph system on spark*, spark) to provide a suitable platform for parallel execution of algorithms. Equipped with great features such as in-memory processing and powerful libraries such as GraphX (*Graphx: A resilient distributed graph system on spark*, spark), Apache Spark can serve as a powerful tool for implementing and executing programs on large graphs. For faster execution, this platform places the needed data on the main memory rather than reading from disk every time. In this study, we used Apache Spark for parallel execution of RWRHN. The feature that makes Apache Spark perfect for reducing algorithm run time when inputs are massive is the ability to partition the input data into several segments then process all segments together with parallel runs of the algorithm.

#### 3.2.1 Parallel computation of ranking with Apache Spark

For parallel execution of RWRHN, it was divided into several parallel steps:

1. each node in the M matrix was assigned with a unique key. Then, for each node, a (*Key, Value*) pair containing the ID of the node and its neighbor and their weight was created and stored. This pair was in the format of (*<u>id</u>*_*<u>2</u>*_, *(id*_*1*_, *weight*)), which means the node *id*_*1*_ resembles the node *id*_*2*_ with the weight of *weight*. The above key/value pair was then partitioned by the key (i.e. *id*_*2*_) using a hash function and the Apache Spark groupByKey command (*Graphx: A resilient distributed graph system on spark*, spark).
2. Another key/value pair called *P(0)* containing the IDs of nodes and their initial rank (*<u>id</u>, rank*) was created and stored and then partitioned with the same method. In this case, the key/value pairs (*<u>id</u>*_*<u>2</u>*_, *(id*_*1*_, *weight*)) and (*<u>id</u>, rank*) which had the same keys were placed in the same partitions.
3. In RWRH, each node receives its rank from its directly linked neighbors. Therefore, it was necessary to find and rank neighboring nodes at each step. To compute the rank of *id*_*1*_, each key/value pairs (*<u>id</u>*_*<u>2</u>*_, *(id*_*1*_, *weight*)) and (*<u>id</u>, rank*) were joined together (provided that *id* = *id*_*2*_) to produce an output in the form of ((*id*_*1*_, *weight*), *Rank*). Since partitioning procedures performed in steps 1 and 2 were by *id* and *id*_*2*_, all of the above data fell in the same partitions and there was no shuffle to slow down the algorithm and add overhead [29]. Finally, at each step of the algorithm, the node *id*_*1*_ received a score from all of its neighboring nodes (i.e. *id*) relative to the corresponding *Rank* value. Since data were initially divided into multiple segments, the Rank value for the set of nodes that were broken into different partitions could be calculated in parallel runs. In other words, breaking the M matrix into multiple partitions allowed the processing cores to compute the rank of a set of nodes simultaneously rather than just one node at a time. For parallel execution of RWRHN, we used the Apache Spark platform implemented on a single computer. In the cases where one machine is not enough to reach a desirable execution speed, the same solution can be expanded by implementing Apache Spark on a group of computers as a processing cluster.

## 4. Parameter setting for the membership functions of type-II fuzzy voter scheme

The type-II fuzzy voter scheme was constructed by the use of the small, medium, and large functions for fuzzification and Vhigh, High, Medium, Low, and Vlow functions for the computation of the agreeability of the sources. Among the several membership functions that can be used for this scheme (e.g. triangular, trapezoidal, etc.), Gaussian functions were chosen because of their excellent accuracy. Since the differences between sources were in the range of (0, 1), the parameters of membership functions were set according to the instructions given in (Mehranfar et al., 2017). This setting is shown in Table.2 and Table.3.

**Table.2.** Parameter values for fuzzification (Mehranfar et al., 2017).

| Spread of lower membership function | Mean of lower membership function |  |  | Spread of upper membership function | Mean of upper membership Function |  |  |
| --- | --- | --- | --- | --- | --- | --- | --- |
| 0.12 | 1 | 0.5 | 0 | 0.16 | 1 | 0.5 | 0 |

**Table.3.** Parameter value for agreeability (Mehranfar et al., 2017).

| Spread of lower membership function | Mean of lower membership function |  |  |  |  | Spread of upper membership function | Mean of upper membership function |  |  |  |  |
| --- | --- | --- | --- | --- | --- | --- | --- | --- | --- | --- | --- |
| 0.06 | 1 | 0.75 | 0.5 | 0.25 | 0 | 0.09 | 1 | 0.75 | 0.5 | 0.25 | 0 |

## 5. Results

This section examines the validity and performance of RWRHN-FF and compares its accuracy with two other methods. Since the proposed method involves using the fuzzy voter scheme to integrate different genome data sources, we also compared the performance of source integration with the conventional averaging method and with the fuzzy voter method. We also used RWRHN-FF to predict new genes for prostate, breast, gastric, and colon cancers and then compared its running time on Apache Spark with the sequential execution. The results of these tests are provided below.

### 5.1 Evaluation of RWRHN-FF

So far, researchers have identified the genes responsible for some diseases, but our knowledge in this area is incomplete and can benefit from further research in this regard. In this study, we used a set of genes that are known to be associated with certain diseases to check the validity of the results of RWEHN-FF using leave-one-out cross-validation (linkage interval and ab initio methods (R. Jiang, Gan, & He, 2011; Y. Li & Patra, 2010)). In the first test (linkage interval), the link between one of the causative genes was removed, other known genes and the disease and its neighbors were designated as seed, and the algorithm was executed to determine whether it can discover the link. The discovery was considered successful if the causative gene was ranked between 1 and 50 by the algorithm. This process was repeated for all known links of the diseases (one link at a time). The second test (ab initio) was similar to the first, except that all genes that are known to be associated with the disease were removed at once. For this test, success was defined as one of the causative genes being placed at the top rank. Having the number of tests and the of successful detections, the precision was calculated accordingly. This measure was defined as the number of successes divided by the total number of tests performed. The results of these two tests for different genes and diseases are presented in Table.4. As can be seen, RWRHN-FF achieved a precision of 55%, while the RWRHN-RE (Luo & Liang, 2015) and RWRH (Y. Li & Patra, 2010) showed a precision of 28% and 23%, respectively. The cause of this superior precision is the strengthening of the disease-disease similarity network and the use of more sources in the construction of gene-gene similarity network. In other words, the constructed heterogeneous network contains more detailed and accurate information than the heterogeneous networks of the rival methods. The ab initio test will result in greater success when having a stronger disease-disease network. In other words, to discover the genes responsible for a specific disease, just the neighbors of that disease will be used. Next, we compared the performance of the proposed algorithm, RWRHN, and RWRHN-RE based on the area under the receiver operating characteristic curve (AUC). We also designed a test to measure the effect of using the type-II fuzzy voter scheme to integrate bio-data source for building reliable gene-gene networks. In this test, the AUC of RWRHN-FF was compared with the AUC of RWRHN in which gene-gene networks were combined using the average of interaction weights. Here, the latter method is referred to as RWRHN-Normal Average (RWRHN-NA).

**Table.4.**
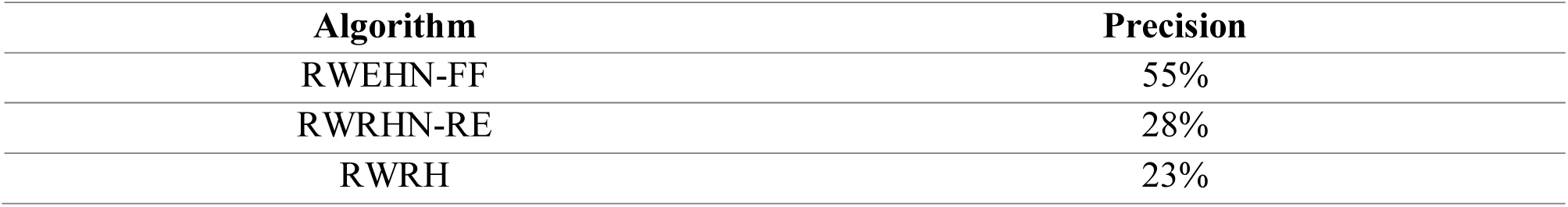
The performance of each method on the Precision.

To calculate the AUC, the sensitivity was plotted against 1-Specificity. Here, the sensitivity is the ratio of disease-causing genes that have been correctly identified by meeting the threshold and 1-Specificity is the ratio of non-causative genes that have been identified as causative genes incorrectly. Fig.6 and Fig.7 illustrate the AUC obtained for RWRHN-NA, RWRHN, RWHN-FF, and RWHN-RE, which are 0.7919, 0.9602, 0.7812 and 0.8667, respectively (presented in Table.5). It can be seen that RWRHN-FF has achieved better precision than other methods; an achievement that must be attributed to its ability to use different genomic sources in the construction of the gene-gene network, to utilize type-II fuzzy voter scheme to reduce the impact of false interactions during source integration, and to use four different disease sources to build disease-disease and disease-gene networks. In other words, because of the use of type-II fuzzy voter scheme, false positives (false interactions) have been given lower weights and have had a reduced impact on the ranking of disease-causing genes. Therefore, compared to RWRHN-NA, RWHN, and RWHN-RE, the proposed method misidentifies a fewer number of non-causative genes as disease-causing ones, which results in better AUC and higher precision. The comparison of RWRHN-NA and RWRHN-RE shows that using more data sources does not necessarily lead to better results and requires a suitable method for integrating data and reducing the effect of false positives.

**Table.5.** The performance of each method on the AUC metric.

| Algorithm | AUC |
| --- | --- |
| RWEHN-FF | 0.9602 |
| RWRHN-RE | 0.8667 |
| RWRH-NA | 0.7919 |
| RWRHN | 0.7812 |

**Fig.6.**
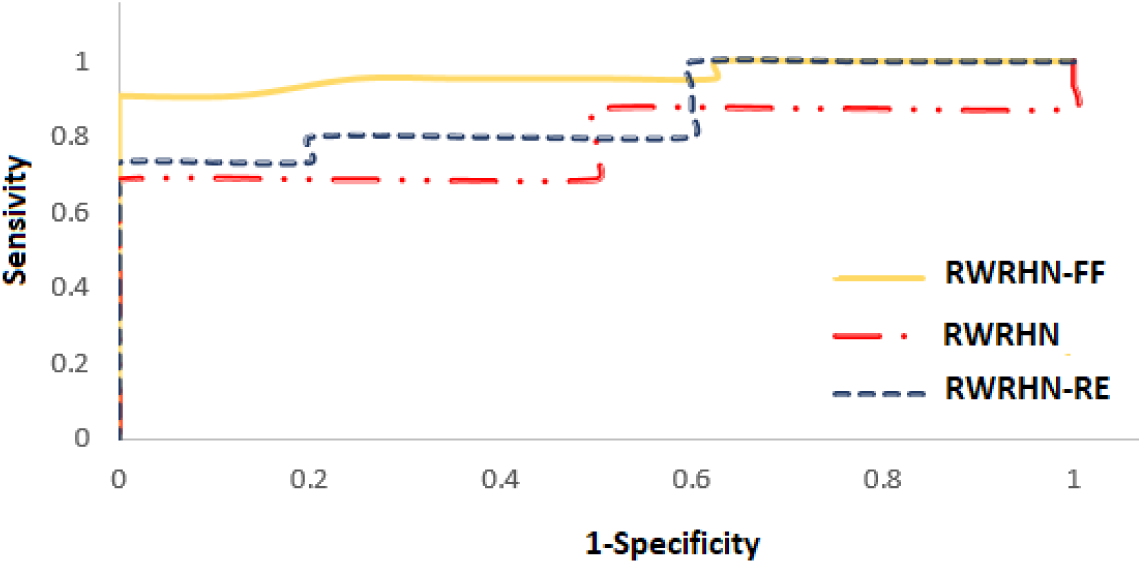
Performance comparison of our method (RWRHN-FF) by RWRHN and RWRHN-RE.

**Fig.7.**
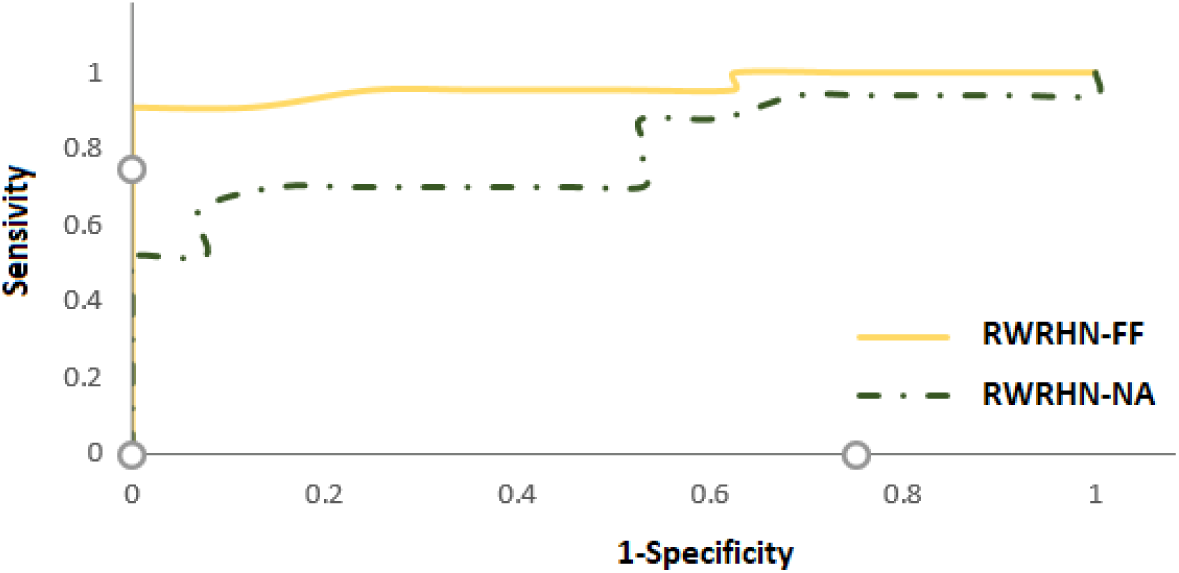
Performance comparison of our method (RWRHN-FF) by RWRHN-NA.

### 5.2 Validity : Prediction of new disease-causing genes with RWRHN-FF

The validity and accuracy of the proposed algorithm can be verified by investigating the causative genes of several diseases. For this purpose, we used the proposed algorithm to identify the genes of breast, gastric, colon, and prostate cancers. The first ten genes predicted by RWRHN-FF for these diseases are listed in Table.6.

For prostate cancer, for example, the genes DMXL1 and RNASEL (Pontén, Jirström, & Uhlén, 2008; Safran et al., 2010) were among the known genes and were present in the input (seed) data. The algorithm predicted eight new genes, among which TP53BP2, SLC12A6, SST, ITGA2, ERBB3, and 4ERBB were correctly detected (Pontén et al., 2008; Safran et al., 2010). For colon cancer, six genes including SMAD2, ARRB2, SMAD3, TBL1XR1, ASS1, and NGB were already present in the heterogeneous network and the algorithm correctly predicted four other genes namely TBL1Y, TBL1X, PAFAH1B, and ASPP1(Pontén et al., 2008; Safran et al., 2010).

### 5.3 Comparison of run time

This section compares the run time of RWRHN-FF in parallel and sequential execution. RWRHN is one of the best-known algorithms for discovering disease-causing genes by the analysis of bio-data networks, but it involves a large number of loops and iterations, which make it time-consuming. For large matrices, RWRHN has a time complexity of O(n^3^) (Symeonidis, Iakovidou, Mantas, & Manolopoulos, 2013). Therefore, in this study, we used Apache Spark as a platform for parallel execution of this algorithm.

For this analysis, a comparison was made between the run time of RWRHN in parallel execution mode and in advanced execution mode (RWRHNFF-Advanced) and the run time of the same algorithm in the two following modes for data of different volumes:

- Sequential execution on a non-Spark platform (Se-RWRHN)

- Non-advanced execution on the Spark platform (RWRHN-NonAdvanced)

In the first mode, the algorithm was executed sequentially on three datasets using Python. In the second mode, RWRHN was executed on the same datasets on the Spark platform but with the non-advanced settings.

In other words, in Se-RWRHN, the ranks of nodes were computed one node at a time. In RWRHN-NonAdvanced, these ranks were computed in parallel; however, in each loop for computing the rank of neighboring nodes, although the key/value pairs (<u>id_2_</u>, (id_1_, weight)) and (<u>id</u>, rank) were joined together, they need shuffling before being joined. In other words, RWRHN-NonAdvanced does not use the hash function for partitioning. In contrast, RWRHNFF-Advanced uses the Spark platform and involves partitioning the key/value pairs (<u>id_2_</u>, (id_1_, weight)) and (<u>id</u>, rank) with the hash function before the rank calculation, which reduces overhead and allows us to avoid shuffle.

Before comparing the run times, the algorithm was executed in Se-RWRHN and RWRHN-NonAdvanced modes to find the causative genes for prostate and breast cancers. In Table 6, the results of Se-RWRHN and RWRHN-NonAdvanced are compared with the results of RWRHNFF-Advanced. As can be seen, the results obtained from these modes of execution are identical.

**Table.6.** Top-10 predicted causal genes of 4 diseases.

| Prostate cancer | Breast cancer | Colon cancer | Gastric cancer |
| --- | --- | --- | --- |
| DMXL1 | BRCA2 | ASPP1 | EGFR |
| TP53BP2 | ATAD5 | SMAD2 | ERBB3 |
| RNASEL | EPC2 | ARRB2 | CD47 |
| SLC12A6 | TOP2A | SMAD3 | VEGFB |
| SST | DHFR | TBL1XR1 | EGR3 |
| ITGA2 | FUS | ASS1 | ERBB4 |
| ERBB3 | RNF19A | NGB | HER2 |
| MYCN | IL1RL1 | PAFAH1B1 | LUM |
| UBL5 | RAD51 | TBL1Y | VEGFA |
| ERBB4 | PIK3CA | TBL1X | DCN |

As also illustrated in Fig.8, when epsilon (convergence condition) was set to 10^−6^, the run times of the compared algorithms for heterogeneous networks of different volumes were as follows. For a heterogeneous network with a size of 1300 MB, RWRHNFF-Advanced convergeed after 4 minutes and RWRHN-NonAdvanced and Se-RWRHN achieved convergence after 11 and 21 minutes, respectively.

**Fig.8.**
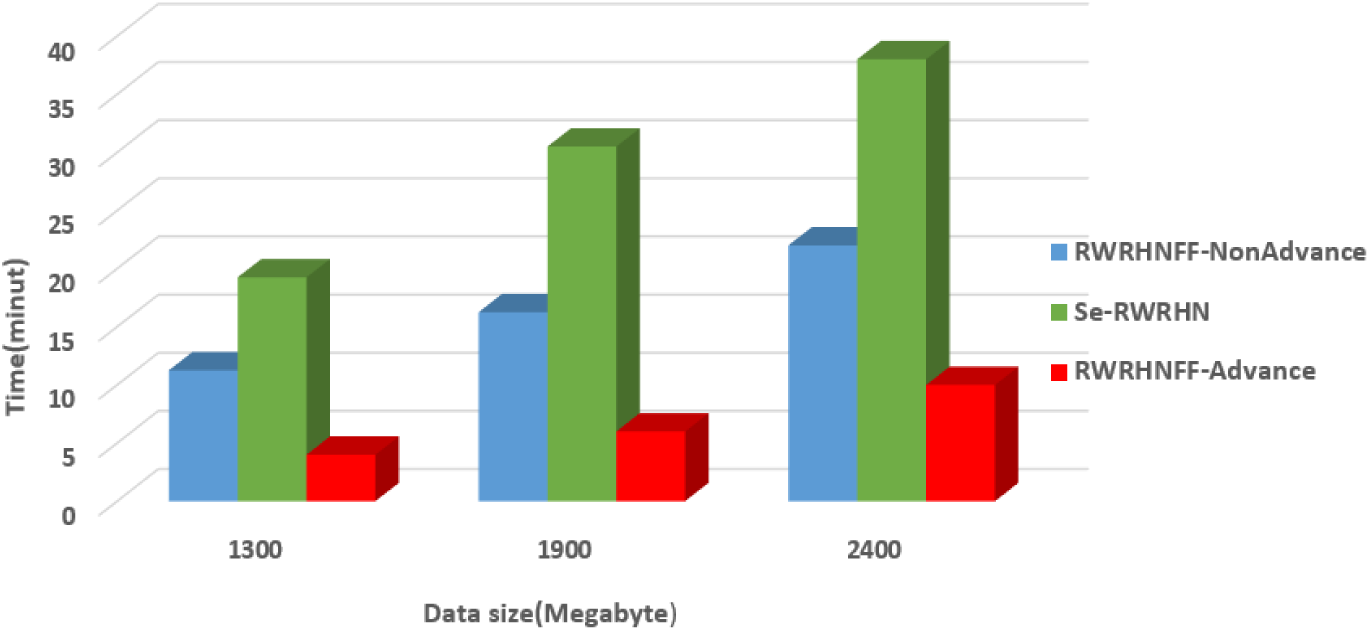
Runtime comparison of parallel execution (RWRHNFF-Advanced and RWRHN-NonAdvanced) against sequential execution (Se-RWRHN).

For a heterogeneous network with a size of 1900 MB, RWRHNFF-Advanced reached convergence after 6 minutes, but RWRHN-NonAdvanced and Se-RWRHN took respectively 16 and 30 minutes to achieve convergence.

For a heterogeneous network with a size of 2400 MB, RWRHNFF-Advanced reached convergence after 10 minutes, but RWRHN-NonAdvanced and Se-RWRHN needed respectively 22 and 38 minutes to achieve convergence.

The above results demonstrate that the parallel execution of RWRHNFF-Advanced on the Spark platform results in faster convergence than its sequential execution (Se-RWRHN). This is primarily because of the partitioning of the heterogeneous network and parallel computation of ranks. In other words, RWRHNFF-Advanced computes the scores of multiple nodes simultaneously instead of one node at a time, as is the case in Se-RWRHN (non-parallel execution). As the results suggest, because of the presence of shuffle procedure in the join operation of non-advanced-RWRHN-SPARK, it has a slower convergence than RWRHNFF-Advanced, which avoids this procedure.

## 6. Conclusion

One of the major missions of bioinformatics is to discover the genes associated with the incidence and development of diseases. Given the importance of this issue, researchers have proposed several approaches to the identification of disease-causing genes. Among the strongest of these approaches is the network-based approach, which typically involves combining gene-gene or protein-protein interaction networks with disease-disease networks. The main problem in working with biomedical networks is that they contain false positives (false interactions) and missing information. However, this issue can be addressed by adding extra bio-data sources to construct new networks or estimating the presence of interactions between each pair of genes or proteins to reduce the impact of experimental errors. Since the use of multiple sources can be effective in reducing the impact of false positives and lack in some sources, we used four sources to reach a more reliable data network. However, having multiple sources does not necessarily lead to a better network and poor data integration can even have a negative impact on the results. In this study, sources were integrated using the type-II fuzzy voter scheme, which was shown to outperform the alternative methods in terms of AUC and Precision. This effect can be attributed to the precise weighting of interactions in this method. Since the use of too many sources in the RWRHN algorithm for the ranking and discovery of causative genes can have a negative impact on its run time, we resolved this issue by using Apache Spark as a platform for parallel execution of this algorithm, which resulted in significantly shorter runtime than sequential execution under similar conditions.

The results of this study demonstrate the possibility of using more data sources to reduce the errors in the existing networks and construct more reliable ones. Since the range and number of membership functions of the algorithm can also influence the final weighting, further examination of the impacts of changes in these functions may prove useful for progress in this area.

## References

Adie, E. A., Adams, R. R., Evans, K. L., Porteous, D. J., & Pickard, B. S. (2005). Speeding disease gene discovery by sequence based candidate prioritization. BMC bioinformatics, 6(1), 55.

Altschul, S. F., Gish, W., Miller, W., Myers, E. W., Lipman, D. J., & (1990). Basic local alignment search tool. Journal of molecular biology, 215(3), 403–410.

Ashkenazy, H., Erez, E., Martz, E., Pupko, T., & Ben-Tal, N. (2010). ConSurf 2010: calculating evolutionary conservation in sequence and structure of proteins and nucleic acids. Nucleic acids research, 38(Suppl_2), W529–W533.

Bateman, A., Coin, L., Durbin, R., Finn, R. D., Hollich, V., Griffiths□Jones, S., … Sonnhammer, E. L. (2011). The Pfam protein families database. Nucleic acids research, 40(D1), D290–D301.

Bock, G. R., & Goode, J. A. (2002). The KEGG database. Paper presented at the ‘In Silico’Simulation of Biological Processes: Novartis Foundation Symposium 247.

Dezső, Z., Nikolsky, Y., Nikolskaya, T., Miller, J., Cherba, D., Webb, C., & Bugrim, A. (2009). Identifying disease-specific genes based on their topological significance in protein networks. BMC systems biology, 3(1), 36.

Graphx: A resilient distributed graph system on spark. (spark). Paper presented at the First International Workshop on Graph Data Management Experiences and Systems.

Ideker, T., & Sharan, R. (2008). Protein networks in disease. Genome research, 18(4), 644–652.

Jaccard, P. (1908). Nouvelles recherches sur la distribution florale. Bull. Soc. Vaud. Sci. Nat., 44, 223–270.

Jiang, R., Gan, M., & He, P. (2011). Constructing a gene semantic similarity network for the inference of disease genes. Paper presented at the BMC systems biology.

Jiang, X., Zhang, H., Quan, X., Liu, Z., & Yin, Y. (2017). Disease-related gene module detection based on a multi-label propagation clustering algorithm. PloS one, 12(5), e0178006.

Karnik, N. N., & Mendel, J. M. (1998). Type-2 fuzzy logic systems: type-reduction. Paper presented at the SMC’98 Conference Proceedings. 1998 IEEE International Conference on Systems, Man, and Cybernetics (Cat. No. 98CH36218).

Karnik, N. N., & Mendel, J. M. (2011). Interval type-2 fuzzy voter design for fault tolerant systems. Information Sciences, 181(14), 2933–2950.

Köhler, S., Doelken, S. C., Mungall, C. J., Bauer, S., Firth, H. V., Bailleul-Forestier, I., … Campbell, J. (2013). The Human Phenotype Ontology project: linking molecular biology and disease through phenotype data. Nucleic acids research, 42(D1), D966–D974.

Le, D.-H., & Dang, V.-T. (2016). Ontology-based disease similarity network for disease gene prediction. Vietnam Journal of Computer Science, 3(3), 197–205.

Lee, I., Blom, U. M., Wang, P. I., Shim, J. E., & Marcotte, E. M. (2011). Prioritizing candidate disease genes by network-based boosting of genome-wide association data. Genome research, gr. 118992.118110.

Li, J., Gong, B., Chen, X., Liu, T., Wu, C., Zhang, F., … Li, X. (2011). DOSim: an R package for similarity between diseases based on disease ontology. BMC bioinformatics, 12(1), 266.

Li, Y., & Patra, J. C. (2010). Genome-wide inferring gene–phenotype relationship by walking on the heterogeneous network. Bioinformatics, 26(9), 1219–1224.

Linda, O., & Manic, M. (2011). Interval type-2 fuzzy voter design for fault tolerant systems. Information Sciences, 181(14), 2933–2950.

Liu, X., Yang, Z., Lin, H., Simmons, M., & Lu, Z. (2017). DIGNiFI: Discovering causative genes for orphan diseases using protein-protein interaction networks. BMC systems biology, 11(3), 23.

Luo, J., & Liang, S. (2015). Prioritization of potential candidate disease genes by topological similarity of protein–protein interaction network and phenotype data. Journal of biomedical informatics, 53, 229–236.

Mehranfar, A., Ghadiri, N., Kouhsar, M., & Golshani, A. (2017). A Type-2 fuzzy data fusion approach for building reliable weighted protein interaction networks with application in protein complex detection. Computers in biology and medicine, 88, 18–31.

Montañez, G., & Cho, Y.-R. (2013). Predicting False Positives of Protein-Protein Interaction Data by Semantic Similarity Measures §. Current Bioinformatics, 8(3), 339–346.

Obayashi, T., & Kinoshita, K. (2010). COXPRESdb: a database to compare gene coexpression in seven model animals. Nucleic acids research, 39(Suppl_1), D1016–D1022.

OMOM. (2014). OMIM. org: Online Mendelian Inheritance in Man (OMIM®), an online catalog of human genes and genetic disorders. Nucleic acids research, 43(D1), D789–D798.

Osborne, J. D., Flatow, J., Holko, M., Lin, S. M., Kibbe, W. A., Zhu, L. J., … Chisholm, R. L. (2009). Annotating the human genome with Disease Ontology. BMC genomics, 10(1), S6.

Oti, M., & Brunner, H. G. (2007). The modular nature of genetic diseases. Clinical genetics, 71(1), 1–11.

Peri, S., Navarro, J. D., Amanchy, R., Kristiansen, T. Z., Jonnalagadda, C. K., Surendranath, V., … Gronborg, M. (2003). Development of human protein reference database as an initial platform for approaching systems biology in humans. Genome research, 13(10), 2363–2371.

Pontén, F., Jirström, K., & Uhlén, M. (2008). The Human Protein Atlas—a tool for pathology. The Journal of Pathology: A Journal of the Pathological Society of Great Britain and Ireland, 216(4), 387–393.

Safran, M., Dalah, I., Alexander, J., Rosen, N., Iny Stein, T., Shmoish, M., … Krug, H. (2010). GeneCards Version 3: the human gene integrator. Database, 2010.

Schlicker, A., Lengauer, T., & Albrecht, M. (2010). Improving disease gene prioritization using the semantic similarity of Gene Ontology terms. Bioinformatics, 26(18), i561–i567.

Shahreza, M. L., Ghadiri, N., Mousavi, S. R., Varshosaz, J., & Green, J. R. (2017). Heter-LP: A heterogeneous label propagation algorithm and its application in drug repositioning. Journal of biomedical informatics, 68, 167–183.

Silberberg, Y., Kupiec, M., & Sharan, R. (2017). GLADIATOR: a global approach for elucidating disease modules. Genome medicine, 9(1), 48.

Stenson, P. D., Mort, M., Ball, E. V., Howells, K., Phillips, A. D., Thomas, N. S., & Cooper, D. N. (2009). The human gene mutation database: 2008 update. Genome medicine, 1(1), 13.

Symeonidis, P., Iakovidou, N., Mantas, N., & Manolopoulos, Y. (2013). From biological to social networks: Link prediction based on multi-way spectral clustering. Data & Knowledge Engineering, 87, 226–242.

Tian, Z., Guo, M., Wang, C., Xing, L., Wang, L., & Zhang, Y. (2017). Constructing an integrated gene similarity network for the identification of disease genes. Journal of biomedical semantics, 8(1), 32.

van Dam, S., Vosa, U., van der Graaf, A., Franke, L., & de Magalhaes, J. P. (2017). Gene co-expression analysis for functional classification and gene–disease predictions. Briefings in bioinformatics.

Van Driel, M. A., Bruggeman, J., Vriend, G., Brunner, H. G., & Leunissen, J. A. (2006). A text-mining analysis of the human phenome. European journal of human genetics, 14(5), 535.

Wang, J. Z., Du, Z., Payattakool, R., Yu, P. S., & Chen, C.-F. (2007). A new method to measure the semantic similarity of GO terms. Bioinformatics, 23(10), 1274–1281.

Wang, X., Gulbahce, N., & Yu, H. (2011). Network-based methods for human disease gene prediction. Briefings in functional genomics, 10(5), 280–293.

Zeng, X., Liao, Y., Liu, Y., & Zou, Q. (2017). Prediction and validation of disease genes using HeteSim Scores. IEEE/ACM Transactions on Computational Biology and Bioinformatics (TCBB), 14(3), 687–695.

Zeng, X., Zhang, X., & Zou, Q. (2015). Integrative approaches for predicting microRNA function and prioritizing disease-related microRNA using biological interaction networks. Briefings in bioinformatics, 17(2), 193–203.

